# Wild gut microbiomes reveal individuals, species, and location as drivers of variation in two critically endangered tropical birds

**DOI:** 10.1101/2021.05.28.446199

**Authors:** Maria S. Costantini, Matthew C.I. Medeiros, Lisa H. Crampton, Floyd A. Reed

**Affiliations:** School of Life Sciences, University of Hawai‘i at Mānoa, Honolulu, HI, USA; Pacific Biosciences Research Center, University of Hawai‘i at Mānoa, Honolulu, HI, USA; Hawai‘i Division of Forestry and Wildlife, Hanapepe, HI 96716, USA; Pacific Cooperative Studies Unit, University of Hawai’i, Honolulu, HI 96822, USA

**Keywords:** Avian microbiome, conservation, Hawaiian honeycreepers, microbial ecology, molecular ecology

## Abstract

The gut microbiome of an animal has a strong influence on the health, fitness, and behavior of its host, and is thus a critical component of the animal itself. Most research in the microbiome field has focused on human populations and commercially important species. However, researchers are now considering the link between endangered species conservation and the microbiome. In Hawai‘i, several threats have caused widespread population declines of Hawaiian honeycreepers (subfamily Carduelinae). These threats, and the environmental changes that result, can have a significant effect on the avian gut microbiome and may even lead to disruption of microbial function. However, no previous study has explored the natural patterns of the gut microbiome of a honeycreeper species in the wild. This project used amplicon-based sequencing of the 16S rRNA gene to characterize the gut microbiome of two critically endangered species of Hawaiian honeycreepers. The two species differed significantly in both alpha and beta diversity. Intraspecific variation of the gut microbiome among individual birds was a major factor. However, small but significant differences also exist between sampling location and sexes. This baseline knowledge will help inform management decisions for these honeycreeper species both in their native habitats and in captivity.

## Introduction

Host-associated microbiota—bacteria, fungi, archaea, protists, and viruses—are often critically important to the healthy physiological functioning of their host (Kropackova *et al*. 2017; Nieves-Ramírez *et al*. 2018). Additionally, different host organs (e.g., gut, skin, vagina) will foster unique microbial communities that possess their own specific function or functions. The vertebrate gut microbiome has been a major focus of microbiome research and is known to provide a suite of functional benefits for the host, including augmenting the immune defense against pathogens, digestion and nutrient acquisition, and processing of dietary toxins (reviewed in McFall-Ngai 2013). Variation in the gut microbiome between hosts can account for differences in host functioning, and understanding the origin of this variation may be crucial to reestablishing optimal host function after a disturbance (e.g., habitat loss, invasion by non-native species, extreme weather; West *et al*. 2019)

There is a significant taxon bias in gut microbiome studies. The vast majority of microbiome research thus far has been on mammals, mostly humans (Grond *et al*. 2018). By comparison, the avian microbiome is poorly described and only recently has there been a push to understand the mechanisms that drive microbiome assembly in wild birds (Colston and Jackson 2016). Microbiome studies on avian species are expected to inherently differ from mammalian species because of the route of initial colonization. Mammals are born through their mother’s vaginal canal, which is rich in microbial species (Dominguez-Bello *et al*. 2010). While there is some evidence that birds may receive maternal transfer of bacteria *in ovo* (Trevelline *et al*. 2018), it is believed that the most of the initial acquisition of microbes happens via the environment (i.e., the nest and parental crop; Grond *et al*. 2017, Chen *et al*. 2020).

Hawaiian honeycreepers (*Passeriformes Drepanididae*) are a particularly interesting avian system for ecological- and evolutionary-based questions because of the substantial degree of phenotypic variation due to adaptive radiation within the lineage (Lovette *et al*. 2002). Despite widespread interest in the group because of this aspect, as well as heightened interest for conservation purposes, no research thus far has focused on the gut microbiomes of Hawaiian honeycreepers or any endemic Hawaiian bird species. Across the lineage, many threats are responsible for population declines; however, avian malaria (*Plasmodium relictum*) carried by the mosquito vector *Culex quinquesfasciatus* is of greatest concern, currently constraining the majority of native forest birds to high elevations (Fortini *et al*. 2015). On the island of Kaua‘i, the situation is particularly dire as climate change is rapidly facilitating the movement of avian malaria into the last stronghold for native forest birds on the island, the Alaka‘i Plateau (Paxton *et al*. 2016). Kaua‘i is also home to two of the most critically endangered honeycreeper species where they only occupy a small (54 km^2^) region of their original range (Fortini *et al*. 2015). The ‘Akikiki (*Oreomystis bairdi*) is estimated to have 468 individuals (231-916, 95% CI) and the ‘Akeke‘e (*Loxops caeruleirostris*) is estimated to have 945 individuals (460-1,547, 95% CI) remaining in the wild (Paxton *et al*. 2016). Both ‘Akikiki and ‘Akeke‘e require immediate and drastic conservation actions; however, unlike many of their contiguous North American counterparts, minimal research has focused on either species and, to some extent, even baseline ecological and life history knowledge is lacking.

The ‘Akikiki and ‘Akeke‘e are both insectivorous species with almost entirely overlapping ranges (Behnke *et al*. 2016). While ‘Akikiki and ‘Akeke‘e are both insectivorous forest birds, they forage in different levels of the canopy and utilize different foraging techniques, potentially resulting in minimally overlapping diets and, thus, minimally overlapping gut microbial communities, as diet is known to strongly affect microbiome composition (Pascoe et al. 2017; Youngblut *et al*. 2019; Teyssier *et al*. 2020). While both species forage mostly within the tree species, ‘ōhi‘a lehua (*Metrosideros polymorpha*), that makes up the majority of the canopy in the native forest bird range, it is believed to be that ‘Akeke‘e are more ‘ōhi‘a-specialists than ‘Akikiki, which are often found foraging in other native tree species (Foster *et al*. 2000; VanderWerf and Roberts 2008). This has been supported by a molecular diet analysis of ‘Akikiki and ‘Akeke‘e that demonstrate that ‘Akikiki have broader diets (M.S. Costantini, unpublished data). It is then plausible that the differences in these foraging niches and diets may translate to the gut microbiome predicting that ‘Akikiki have higher bacterial diversity due to their more generalized diet.

In addition to diet, there are several other factors known to influence microbiome composition in some species both intrinsically (e.g. age, breeding condition, sex, evolutionary history) and extrinsically (e.g. nesting environment, local prey availability, behavioral interactions) (Spor *et al*. 2011; Grond *et al*. 2018). In the context of conservation, many of the threats that are causing the decline of honeycreeper populations (e.g., habitat degradation, altering prey availability, the introduction of novel species) and the responding management actions (e.g. captive breeding) can also affect the host-associated microbiome by altering the available microbial species pool in the environment (Carthey *et al*. 2020).

As both species face greater risks of extinction and subsequent conservation actions are implemented in response, it is imperative that a baseline understanding of their gut microbiomes in the wild is understood. We targeted the bacterial community, specifically, by sequencing the 16S rRNA gene and conducted statistical analyses focused on comparisons of alpha and beta diversity metrics between and within host species and at two different locations (one on the periphery of the receding Kaua‘i forest bird range and one in its core). The goal of this project is to describe the natural patterns in the gut microbiome of ‘Akikiki and ‘Akeke‘e as they relate to species identity, temporal and spatial sampling patterns, and intraspecific variation of the hosts.

## Methods

### Field Sampling

We collected a total of 13 fecal samples from 13 individual ‘Akeke‘e and 41 fecal samples from 36 individual ‘Akikiki between 2017 and 2018 except for one ‘Akikiki sample that was collected during the winter of 2016. Samples were collected from across the Alaka‘i Plateau on the island of Kaua‘i, Hawai‘i, which represents the remaining range of both species in the wild (Fig.1). The main field site of the study, Halepa‘akai, is located on the eastern side of the plateau and contains the highest abundance of both species (Behnke *et al*. 2016). They are also observed, though much less regularly, at the Upper Kawaikōi field site. Our study sites on the plateau range from an elevation of 1,400 m in the east at Halepa‘akai to about 1,200 m in the west at Upper Kawaikōi.

**Figure 1.**
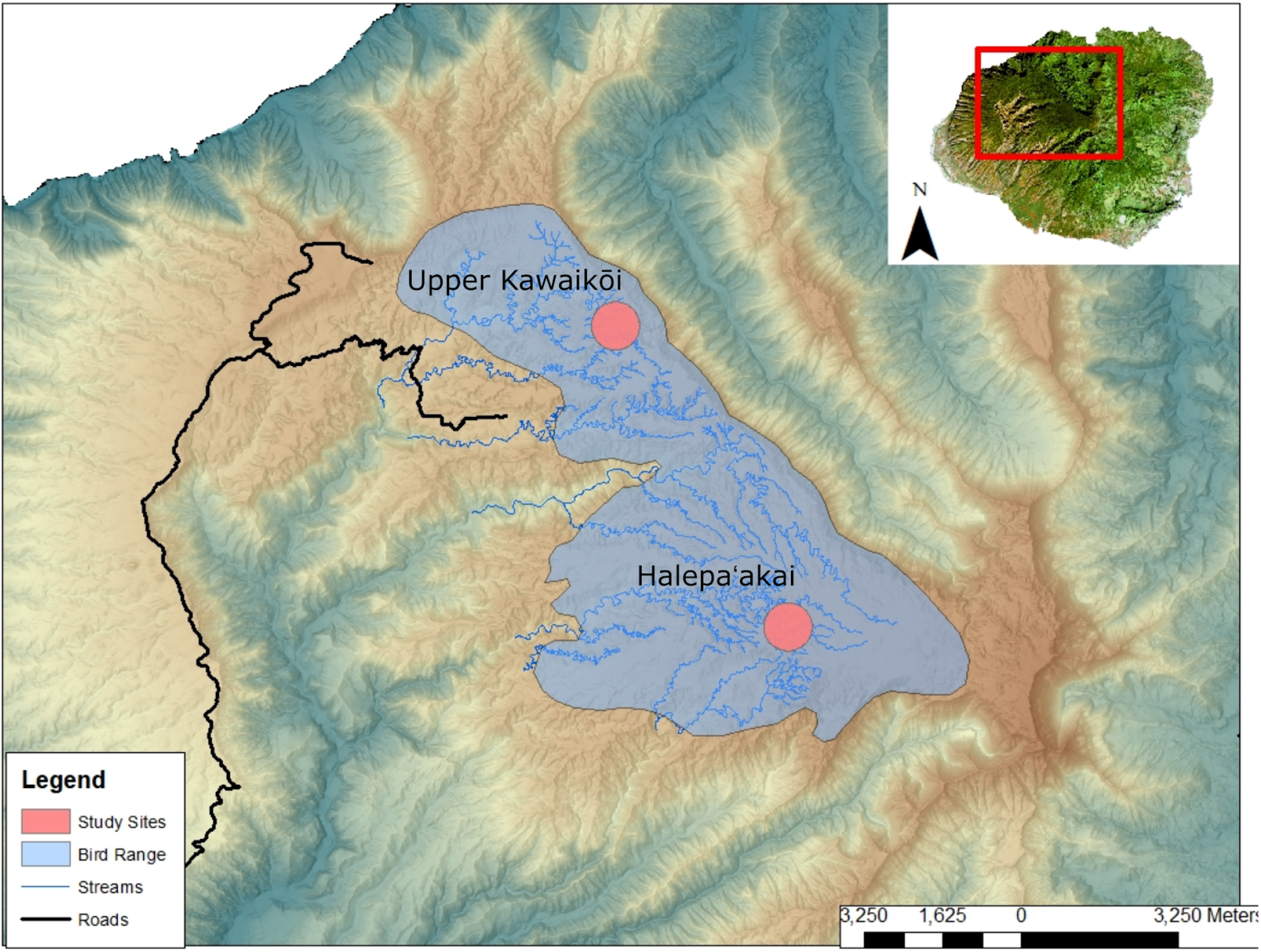
Sampling locations within the remaining endangered forest bird range on the island of Kaua‘i.

To analyze the gut microbiome we used fecal samples as a proxy for the gut as they represent the most accurate view of the colon microbiome; short of sacrificing individuals and harvesting the gastrointestinal tract (Videvall *et al*. 2018). The Kaua‘i Forest Bird Recovery Project (KFBRP) has been opportunistically collecting fecal samples through mist netting and banding since the early 2000’s (under IACUC #08-585-7). Samples were collected year-round, though the majority of samples were taken between January and June due to increased bird activity during the breeding season. Upon capture, birds were placed in sterile cloth bags for no more than 30 minutes. Each bag was only used once for a single bird, and then retired, until it could be washed with bleach and hot water. Fecal samples were directly collected from bags into 1.5 mL Eppendorf tubes containing 100% ethanol and later stored at −20° C for longer term preservation. Upon capture we placed unique aluminum USGS leg bands on each individual, recorded age and sex, and recorded morphometric measurements to track body condition.

### Sequencing

We used a Qiagen DNeasy Powersoil Kit to extract DNA from fecal samples by following the manufacturer’s instructions with modification for arthropod diets (Qiagen, Germantown, MD, USA). To remove ethanol from samples prior to extraction, we conducted two washes of the fecal pellet with RNA/DNA free molecular grade water as described in Grond *et al*. (2014). Extracted DNA was quantified and quality-controlled using the Invitrogen Qubit 4 Fluorometer (ThermoFisher Scientific, Waltham, MA, USA). Sequencing for bacterial taxa was conducted on the V4 region of the bacterial 16S rRNA gene using forward primer 515F (5’-GTGYCAGCMGCCGCGGTAA-3’) and reverse primer 806R (5’-GGACTACNVGGGTWTCTAAT-3’). The PCR cycling conditions adhered to the Earth Microbiome Project protocol and were as follows: an initial denaturing step at 94°C for 3 min; followed by 35 cycles of 94°C for 45 s, 50°C for 60 s, and 72 °C for 90 s, and then a final extension period at 72 °C for 10 min (Gilbert *et al*. 2014). 16S amplicons (PCR products) were index tagged to identify the originating bird sample and location, and sequenced using the Illumina MiSeq platform with the v3 (2 × 300 cycles) reagent kit. All library preparation and sequencing was conducted at the Advanced Studies in Genomics, Proteomics, and Bioinformatics (ASGPB) facility at the University of Hawai‘i at Mānoa (Honolulu, Hawai‘i, USA).

### Data Processing and Statistical Analyses

The University of Hawai‘i at Mānoa‘s C-MAIKI (Center for Microbiome Analysis through Island Knowledge and Investigation) pipeline (Arisdakessian *et al*. 2020) for amplicon-based microbiome analysis was used to process samples from raw reads to amplicon sequence variants (ASVs). ASVs are similar in concept and utility as the more traditionally used operational taxonomic units (OTUs). However, unlike OTUs, ASVs are resolved down to the level of single-nucleotide differences. This allows for finer-scale resolution than the OTU clustering methods (Callahan *et al*. 2017). The C-MAIKI pipeline quality-filtered and denoised sequences in the program DADA2, then aligned, filtered, and annotated sequences in the program MOTHUR using the Silva database (Schloss *et al*. 2009, Callahan *et al*. 2016, https://www.c-maiki.org/). Potential chimeras were removed with VSEARCH (Rognes *et al*. 2016) through MOTHUR. Sequences matched to chloroplasts, archaea, mitochondria, and *Wolbachia* were removed from the dataset.

Statistical analyses processing and data visualization was performed in R using the phyloseq, ggplot2, and vegan packages (R Core Team 2013; Oksanen *et al*. 2007; McMurdie and Holmes 2013). The alpha diversity of each gut microbiome was determined by calculating the Chao1 and Shannon diversity indices and tested for significant differences between groups using generalized linear models (GLM) with sequencing depth included as a covariate. Differences in gut microbiome beta diversity between certain groups (i.e., species, sampling location, sampling season and sex) were visualized using non-metric multidimensioal scaling (NMDS) plots with the Bray-Curtis dissimilarity index. To account for differences in sampling depth, we randomly down-sampled (rarefied) to the same read count per sample (16,068 reads per sample). Significance was then tested by calculating non-parametric, permutational multivariate analyses of variance (PERMANOVA; ‘vegan’: *adonis*) with 10,000 permutations.

## Results

The 16S rRNA sequencing of our samples yielded 2,667,386 quality filtered reads (range: 16,068-88,894 sequences per sample). After preprocessing the data to remove biologically irrelevant taxa and extremely rare taxa, we identified a total of 8,958 ASVs. Rarefaction curves for each individual based on ASV richness indicated that our sequencing depth was sufficient for capturing alpha diversity (Supplementary Fig. 1). We removed sample “369” from analysis, as it did not appear to amplify.

### Interspecific Differences in Gut Microbiota

‘Akikiki and ‘Akeke‘e have distinct microbial communities from one another, as evidenced by minimal overlap between species ellipses when plotted with Bray-Curtis distance measurements (Supplementary Fig. 2; PERMANOVA: R^2^ = 0.10, p = 9.999 x 10^-5^). Despite differences in beta diversity, there were still similarities regarding the individual taxa between the two communities. The gut microbiomes of both ‘Akikiki and ‘Akeke‘e were generally dominated by bacteria from the phyla Proteobacteria, Firmicutes, and Actinobacteria, but the relative abundances of each phylum differed (Fig. 2). Proteobacteria were more dominant in the ‘Akeke‘e microbiome (65.3% average relative abundance) than in that of ‘Akikiki (31.1%). In all but two ‘Akeke‘e, Proteobacteria made up nearly 50% or greater of all phyla in the gut. The other two individuals’ microbiomes were dominated by Firmicutes. In contrast, Proteobacteria was at near equal levels with Actinobacteria (30.2%) in the ‘Akikiki microbiome. In both species Firmicutes were at similar levels and the third most abundant, generally (13% in ‘Akeke‘e and 16.7% in ‘Akikiki). Another notable difference between the two species was the relatively high occurrence and abundance of Cyanobacteria in ‘Akikiki. Cyanobacteria contributed more than 1% of the relative abundance in 87.8% (36/41) of ‘Akikiki samples versus only in 54% (7/13) of ‘Akeke‘e samples. Furthermore, in several ‘Akikiki individuals, Cyanobacteria comprised a relatively high proportion of the total bacterial abundance (5.4%) compared to in ‘Akeke‘e (1.1%).

**Figure 2.**
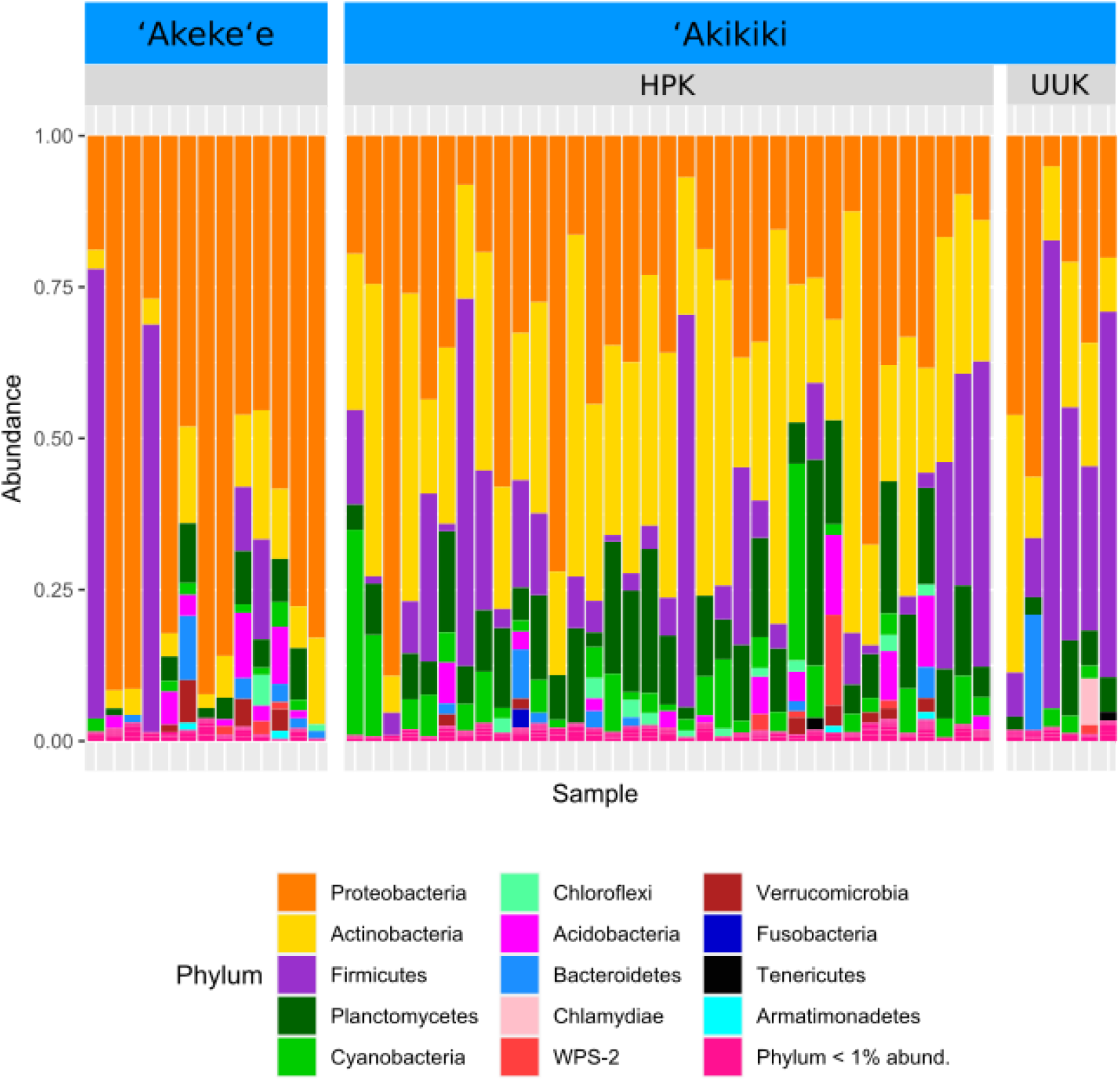
Comparing the relative abundance of bacterial phyla in ‘Akikiki and ‘Akeke‘e. Phyla that make up less than 1% of the read counts of a sample are grouped together in “Phylum <1% abund.”. Also shown is the comparison of bacterial phyla between the two sampling sites for ‘Akikiki. Halepa’akai (HPK) field site represents the core range for the species where occupancy rates are highest and the habitat is considered near pristine native vegetation. Upper Kawaikōi (UUK) is on the fringe of the ‘Akikiki’s present range and has a high density of non-native understory vegetation.

‘Akikiki had a more diverse microbiome than ‘Akeke‘e in terms of the Shannon Index (GLM: p = 0.018; Fig. 3)

**Figure 3.**
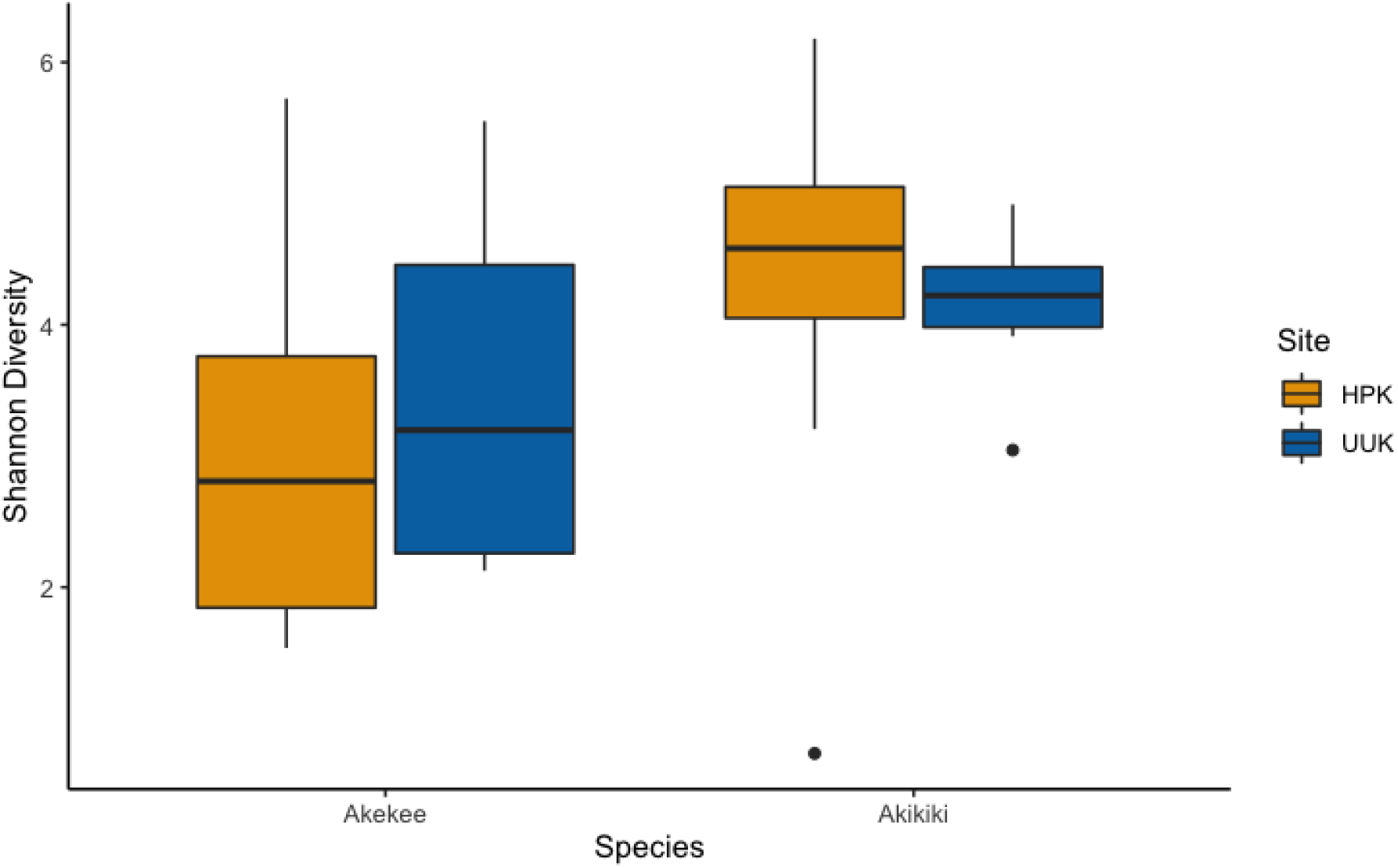
Boxplot of alpha diversity measurements of ‘Akikiki and ‘Akeke‘e at the two sampling locations. Shannon Index estimates species abundance and evenness of ASVs. There was a significant difference between species (GLM: p = 0.018), but not between sites (GLM: p > 0.05).

### Age Differences

To investigate the patterns that drive differences among individuals within a species, we chose to look at only ‘Akikiki samples because we had a more robust sample size. Alpha diversity did not differ between juvenile and adult birds when looking at all pooled samples (GLM: p > 0.05). We used samples from two ‘Akikiki that were sampled as nestlings and resampled as second-year individuals to explore any patterns in longitudinal acquisition of microbiome members. The bacterial community diversity increased in both birds between nestling and second-year stage (Supplementary Fig. 3; this was not tested and is only presented anecdotally because of small sample size, *n* = 2). There was also a notable shift in composition of the most abundant bacterial phyla. The nestlings’ microbiomes were dominated by Actinobacteria, Proteobacteria, and phyla that compromised less than 5% of the relative abundance of all phyla (Supplementary Fig. 4). One of the nestlings also possessed a small proportion of Planctomycetes. Cyanobacteria were present in higher proportions in both second-year individuals, and Firmicutes made up more than 5% of the microbiome in one of the second-year birds.

### Sex Differences

Alpha diversity levels were similar between female and male ‘Akikiki (GLM: p > 0.05), but differed in beta diversity based on Bray-Curtis dissimilarity (PERMANOVA: R^2^ = 0.048, p = 0.032). One bacterial class, Negativicutes, was distinctly different between the two sexes (Fig. 4). Negativicutes were present in only one female, but were found in nine out of the 18 male samples. Other classes, specifically Melainabacteria, Mollicutes, and Phycisphaerae, were uncommon in females but present at moderate levels in males.

**Figure 4.**
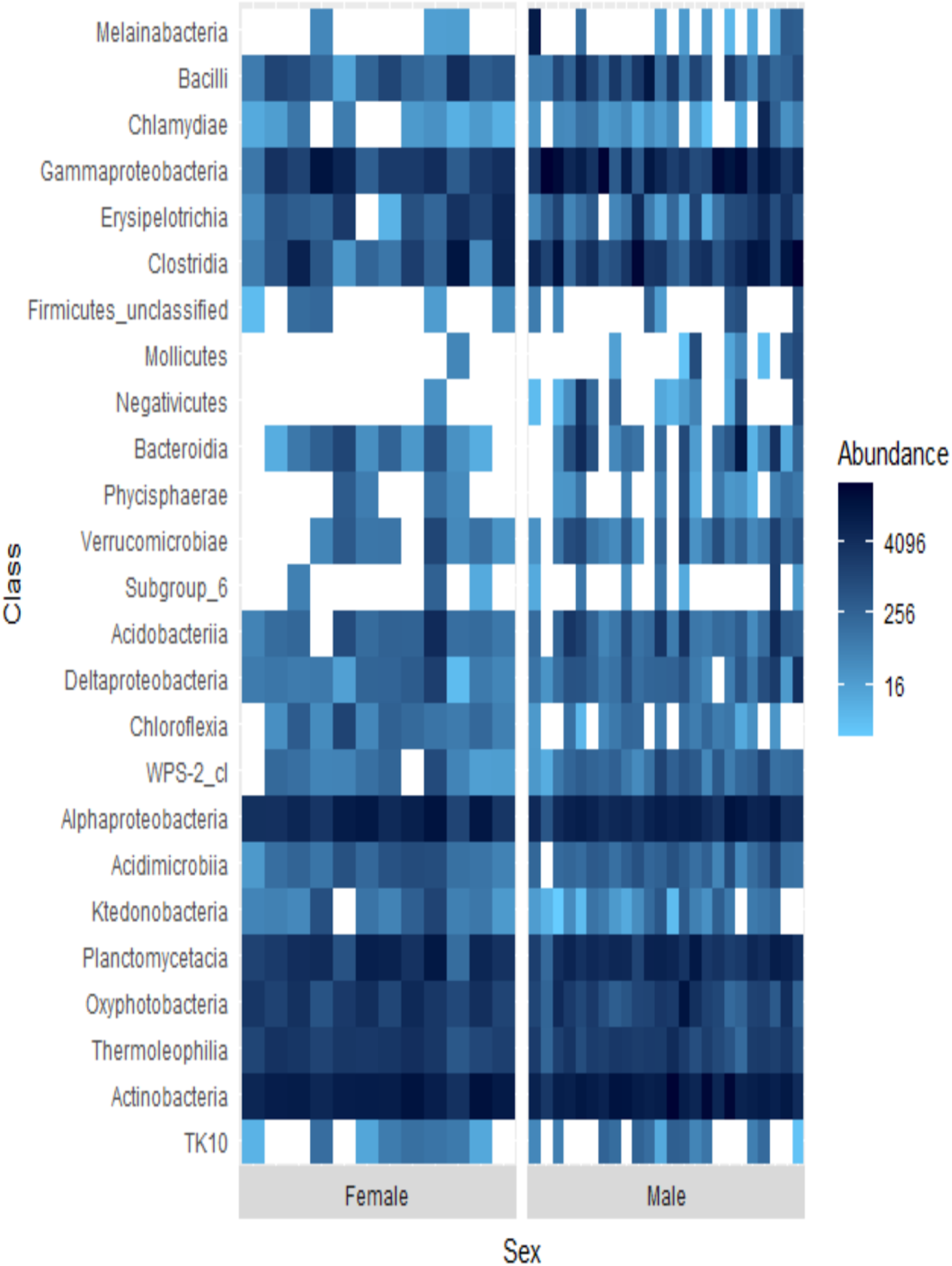
Heatmap showing the relative abundances of the top 25 most abundant bacterial classes in male and female ‘Akikiki. Each box represents an individual sample. Darker blue boxes indicate a greater number of reads of that bacterial class and lighter blue boxes indicate less reads. White boxes indicate that there were no reads of that class in an individual sample.

### Sampling Location Differences

Temporally, we found no difference in the microbiome structure based on sampling season (classified as “Spring/Summer” or “Winter/Fall”; PERMANOVA: R^2^ = 0.036, p = 0.41), but there were spatial differences in the bacterial community structure of ‘Akikiki samples between the two sampling sites (PERMANOVA: R^2^ = 0.064, p = 0.0011). Additionally, there was a shift in the relative abundance of key bacterial phyla between the two sites. Actinobacteria comprised fewer of the total phyla in microbiomes of individuals from the Upper Kawaikōi site (Fig. 2). Cyanobacteria and Planctomycetes, two less dominant phyla, were present in higher proportions of ‘Akikiki microbiomes from Halepa‘akai. Notably, Cyanobacteria was both present in higher proportions within individuals from Halepa‘akai and was present at greater than 1% abundance in 33 of the 35 (94.3%) ‘Akikiki sampled at the location. Only three of the six ‘Akikiki sampled at Upper Kawaikōi possessed Cyanobacteria at the same minimal abundance. There were no differences in Chao1 and Shannon diversity values between the Halepa‘akai and Upper Kawaikōi sites (GLM: p > 0.05; Fig. 3).

## Discussion

The overarching goal of this study was to characterize the wild gut microbiome of two critically endangered Hawaiian honeycreepers and investigate the processes that may influence unique patterns in community composition. Using 16S rRNA amplicon sequencing, we found that the gut microbiomes of ‘Akikiki and ‘Akeke‘e were distinct from one another, and that ‘Akikiki had more diverse microbiomes. This is consistent with the broader foraging and more diverse diet of ‘Akikiki relative to ‘Akeke‘e; supporting the notion that diet is an important driver of microbiome community assembly in these species. A broad look at the similarities within ‘Akikiki and ‘Akeke‘e gut microbiomes reveals that our findings are supported by several wild bird microbiome studies. Wild bird studies generally conclude that Proteobacteria, Firmicutes, and Actinobacteria tend to be the three of the four most dominant phyla in the gut microbiomes (Lewis *et al*. 2017, Kropackova *et al*. 2017, Hird *et al*. 2015, Bodawatta *et al*. 2018, Grond *et al*. 2018). ‘Akeke‘e microbiomes are more dominated by Proteobacteria, while ‘Akikiki are more equally dominated by Proteobacteria and Actinobacteria, and to a lesser extent, Firmicutes. Additionally, both Planctomycetes and Cyanobacteria are more abundant in the ‘Akikiki gut microbiome than in that of ‘Akeke‘e. As ‘Akikiki are thought to have a less specialized diet, the greater proportional spread of bacterial phyla in ‘Akikiki compared to ‘Akeke‘e may be a reflection of this dietary difference. Cyanobacteria, which are photosynthetic prokaryotes, are relatively enriched in the ‘Akikiki microbiome compared to not only ‘Akeke‘e, but to other passerine microbiome studies as well (e.g., Hird *et al*. 2015). A potential explanation for their higher proportional abundance may come from ‘Akikiki foraging ecology. Cyanobacteria are oftentimes the phototrophic partners that live in symbiosis with fungi to make up the complex organisms known as lichen (Nübel *et al*. 1997). ‘Akikiki forage by picking through pieces of moss and lichen to find arthropod prey. Thus, the Cyanobacteria in the ‘Akikiki microbiome could either be obtained directly by inadvertent consumption of lichen or indirectly through arthropod prey that had consumed lichen. Another explanation for the abundance of Cyanobacteria in ‘Akikiki is through colonization from water droplets in the moss in which they forage, as many Cyanobacteria live in association with mosses (Rousk *et al*. 2013).

While inter-specific differences exist in our study, most variation within microbiomes was explained by individual differences. To further investigate potential drivers of individual variation, we looked at environmental and life history characteristics in ‘Akikiki only. We found no difference in alpha or beta diversity in the gut microbiome based on sampling season or age class. However, there is some evidence suggesting that nestlings and adults harbor distinct microbial communities, based on the repeated measures of two individuals (Supplementary Fig. 3 and 4). It was not until both individuals were second-year birds that Cyanobacteria was detected at a notable level. Further investigation into nestling microbiomes will be a key focus for these species moving forward, as disruptions to the microbiome at an early stage is known to lead to lifelong issues (Knutie *et al*. 2017).

We found weak support for a difference between male and females that may be driven by a few key taxa (Fig. 4). Differences in the gut microbiomes between sexes within bird species can be driven by behavior (e.g., differences in foraging ecology) or physiological differences (e.g. impacts of sex hormones; Grond *et al*. 2017). How, and the extent to which, these factors influence microbial community patterns will vary by host species. In the current study we found a higher presence of the bacterial classes Melainabacteria, Mollicutes, Phycisphaerae, and especially, Negativicutes in males than females. Another study on wild birds found an association with Negativcutes and male birds (Liu *et al*. 2020).

Lastly, we explored the possibility that sampling habitat affected the gut microbial community of ‘Akikiki. Bacterial communities differed significantly in beta diversity between Halepa’akai and Upper Kawaikōi within the species. Halepa’akai is considered the last stronghold for the species, where the vast majority of the population exists. Furthermore, it is a relatively pristine, undisturbed forest with little inundation by non-native vegetation (Behnke *et al*. 2016). In contrast, the sub-population at Upper Kawaikōi was recently discovered in 2018 and represents a small fraction of the total population. The habitat is still dominated by ‘ōhi‘a lehua, but the understory has a greater occurrence of non-native and invasive vegetation, including Kahili ginger (*Hedychium gardnerianum*) and strawberry guava (*Psidium cattleyanum*; M.S. Costantini personal observation). Additionally, for an undetermined reason, many of the trunks of ‘ōhi‘a trees in the Upper Kawaikōi site were stripped of moss and lichen (M.S. Costantini personal observation). Here, again, we saw an interesting pattern with Cyanobacteria. The phylum was present at a level of greater than 1% in the gut microbiome of 94.3% of ‘Akikiki samples at Halepa‘akai, but only in 50% of samples from Upper Kawaikōi. This distinction may again be explained by diet or foraging differences between birds at the two sites. A less pristine habitat may result in diminished quality and quantity of prey or differences in the environmental microbial community. It is particularly interesting that Cyanobacteria was enriched in the gut microbiome of birds from the more intact site as a recent study on the microbiome of the American white ibis (*Eudocimus albus*) found that Cyanobacteria significantly decreased in relative abundance with an increase in urban land cover (Murray *et al*. 200). Therefore the presence and abundance of Cyanobacteria in the avian gut microbiome may be an indication of the quality of habitat for certain species. Several studies that have investigated the effect of habitat degradation or land use change in wild animals have found that animals inhabiting degraded or altered habitats have distinctly different microbiomes, often as a result of shifting food availability (reviewed in Trevelline *et al*. 2019). In addition to qualitative changes, degraded habitats often have a reduced alpha diversity. Our results do not show this for ‘Akikiki, but follow up work is necessary with a larger sample size (Fig. 3).

This study represents the first examination of Hawaiian honeycreeper gut microbiomes in the wild. Our goal was to characterize the associated bacterial communities of two critically endangered species to understand the natural patterns and processes in the wild. ‘Akikiki and ‘Akeke‘e will face many conservation challenges in the near future, as 100% of their suitable habitat is predicted to disappear within this century as the parasite that causes avian malaria spreads into higher elevations as a result of climate change (Paxton *et al*. 2016, Fortini *et al*. 2015). Currently, “insurance” populations of both species are being established in captivity; however, several studies on other captive animals have demonstrated a distinct shift in the microbial communities when in captivity (reviewed in Trevelline *et al*. 2019). This disruption can affect the host in numerous ways, including interfering with nutrient acquisition or a weakened immune response to pathogens. This is particularly concerning for Hawaiian honeycreepers, whose major threat in the wild is avian malaria (Paxton *et al*. 2018). While the results from this analysis only provide an exploratory survey of the bacterial members of the gut microbiome of ‘Akikiki and ‘Akeke‘e, it is a necessary first step that must be taken before effective management of the microbiome is possible. Future work will need to focus on determining which microbes are considered symbiotic with the host rather than transient species or environmental contaminants and how the communities are functionally important. Given the critical connection between the gut microbiome and proper physiological functioning of the host, it is imperative that the role of the microbiome be considered as conservation management plans move forward with these species.

## Acknowledgements

This work was funded by Hawai‘i Audubon Society, Watson T. Yoshimoto Fellowship, C-MAIKI Fellowship, and the Kaua‘i Forest Bird Recovery Project. F.A.R’s work is supported by a grant from NIH COBRE P20 GM125508. We thank field technicians and volunteers with the Kaua‘i Forest Bird Recovery Project that helped with banding and sample collection. Particular thanks to J. Hite, T. Winter, and E. Abraham for help in the field. C. Arisdakessian contributed greatly to bioinformatic analysis. N. Yoneishi aided with laboratory work. Bird banding was carried out under federal USGS Bird Banding Laboratory (permit no. 08487). All research was in accordance with the University of Hawai‘i at Mānoa’s Institutional Animal Care and Use Committee (#08-585-7).

---

**Supplementary Figure 1.**
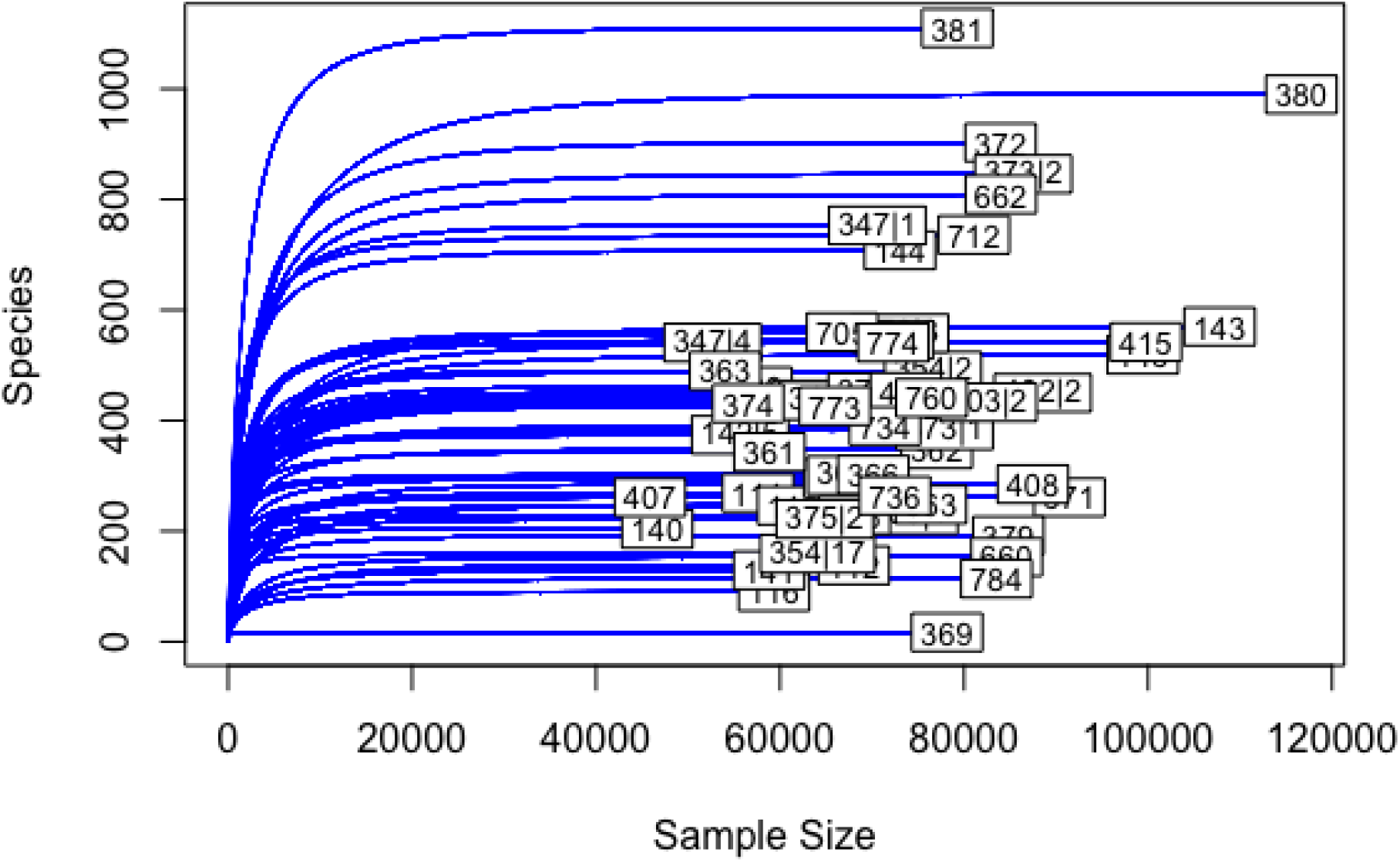
Rarefaction curves for each individual in the study comparing the bacterial ASV richness to sequencing depth.

**Supplementary Figure 2.**
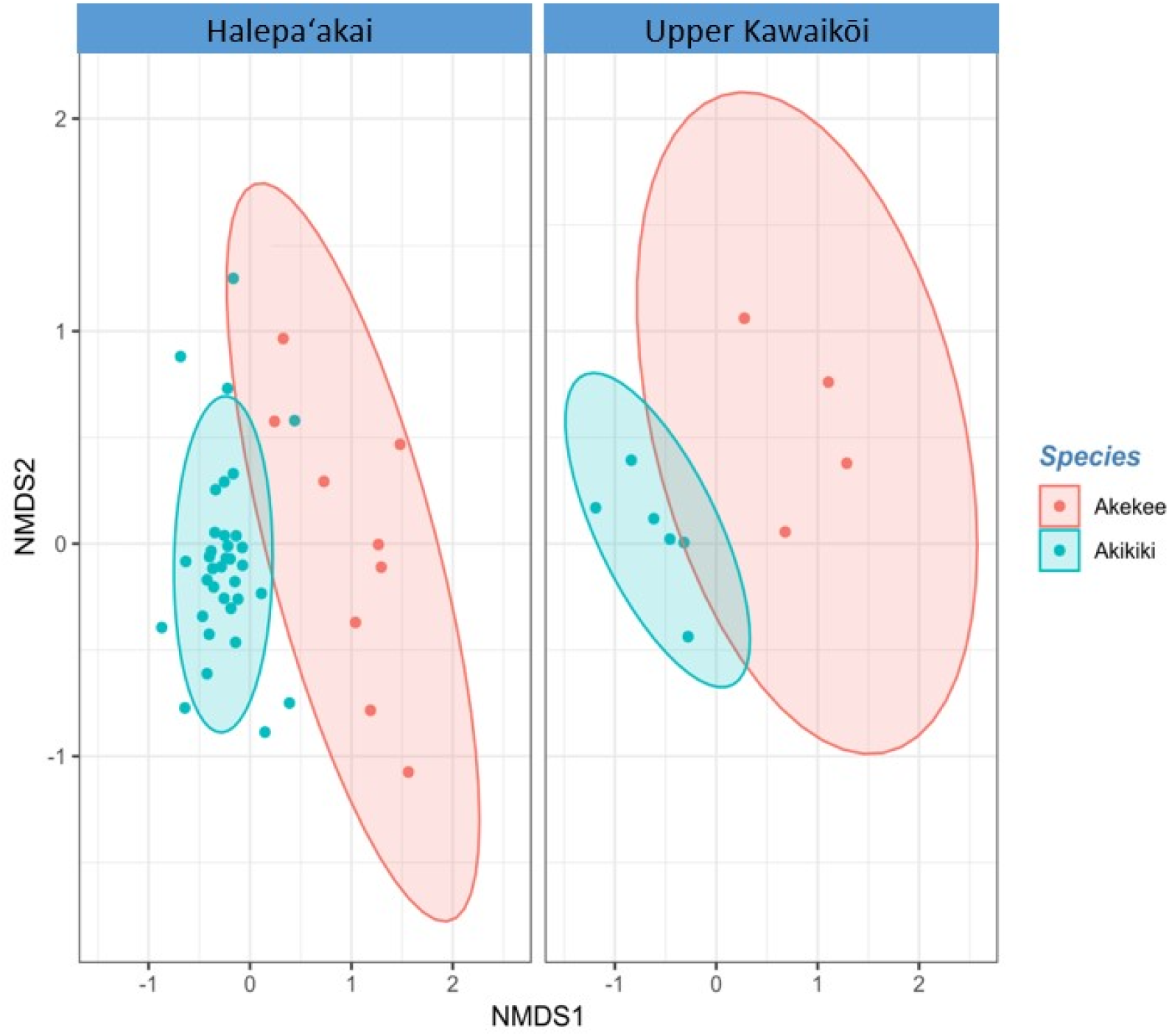
Non-metric multidimensional scaling (NMDS) ordination of Bray-Curtis dissimilarity distances for ‘Akikiki and ‘Akeke‘e. Ellipses represent 90% confidence intervals following a multivariate t-distribution.

**Supplementary Figure 3.**
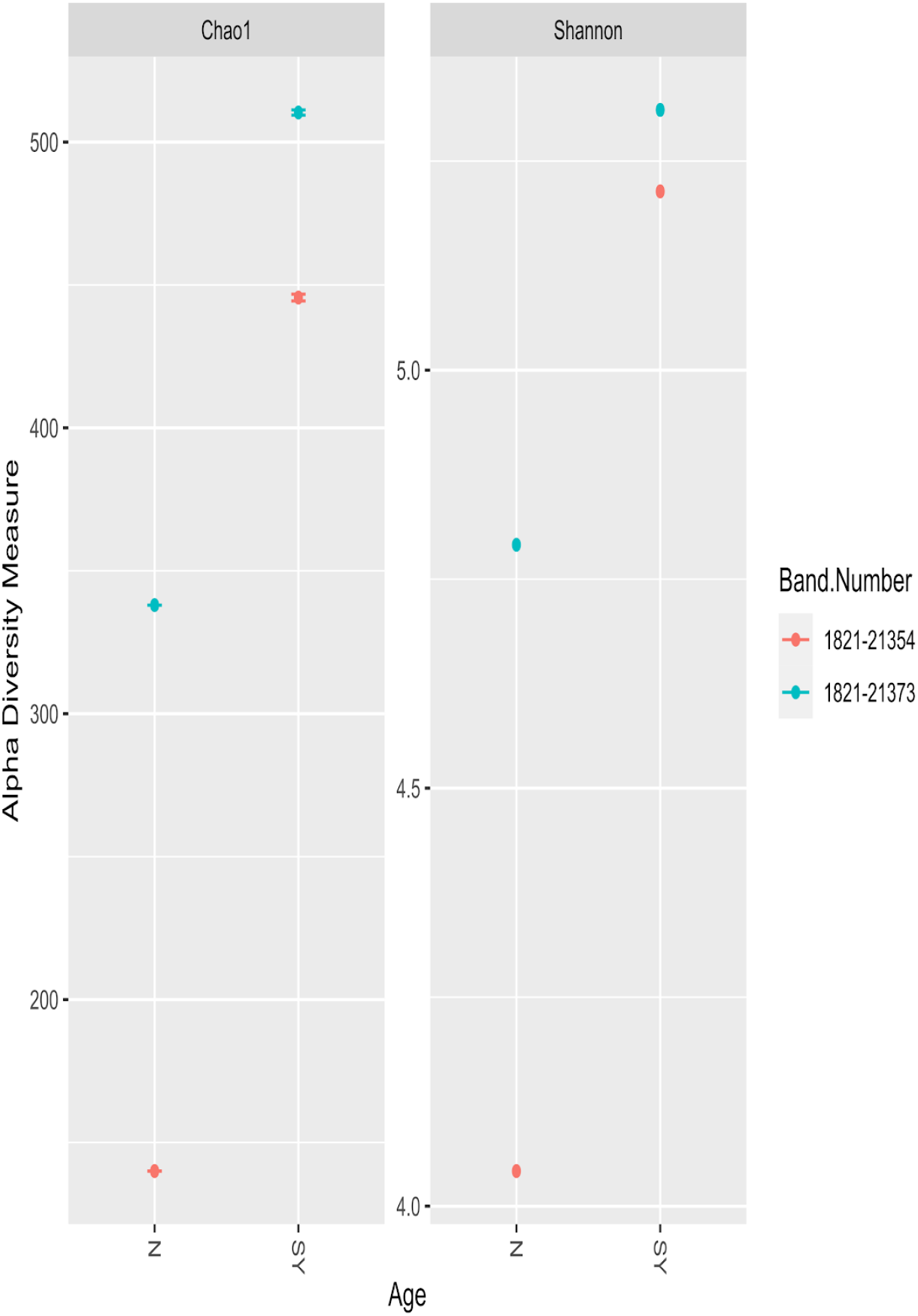
Alpha diversity measurements for two ‘Akikiki individuals as nestlings (N) and second-year (SY) birds. Chao1 values represent nonparametric species richness of ASVs and the Shannon index estimates species abundance and evenness of ASVs. Each dot is one sample.

**Supplementary Figure 4.**
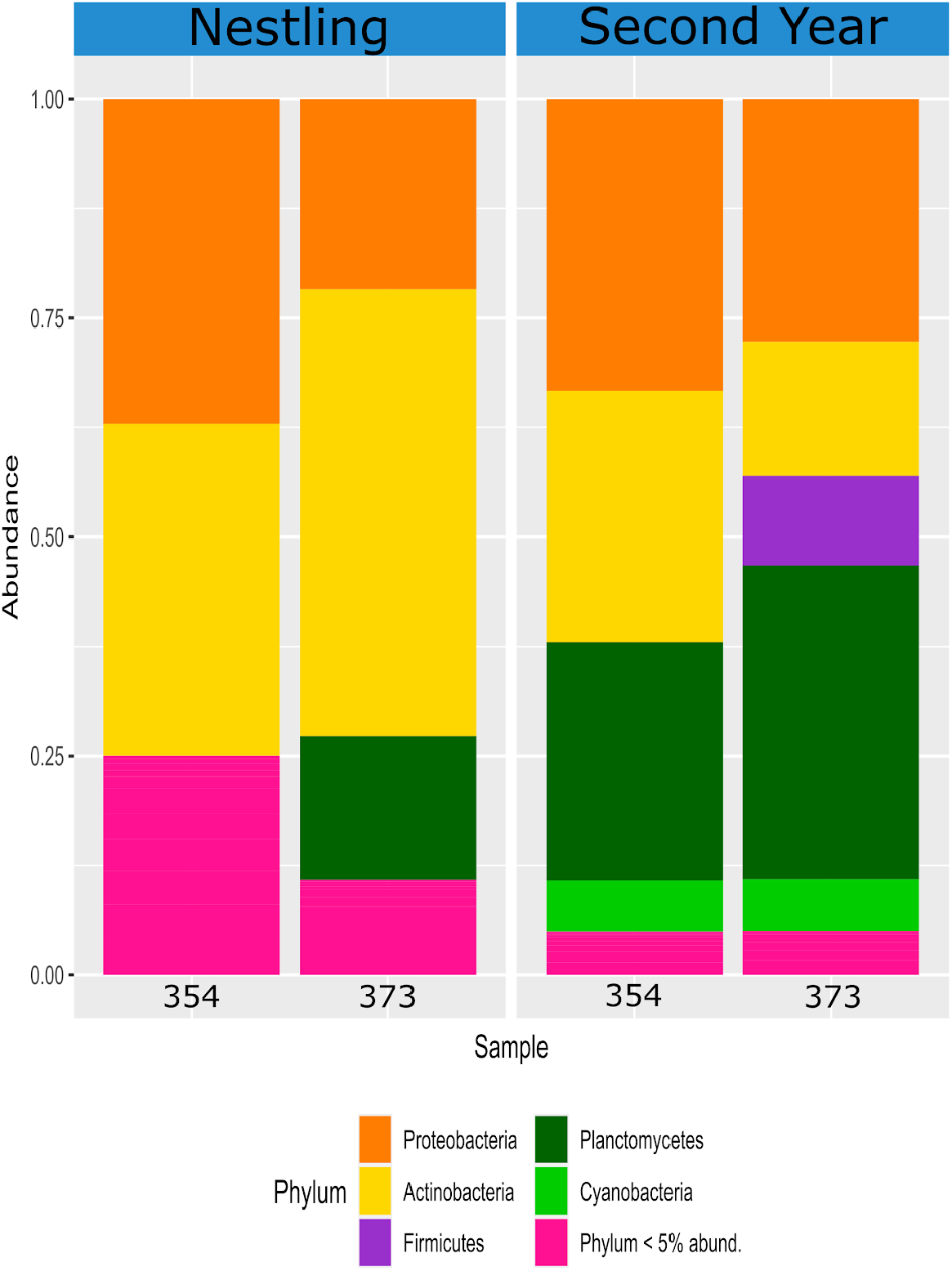
Comparing the relative abundance of bacterial phyla of two ‘Akikiki individuals sampled at different life stages. Phyla that make up less than 5% of the read counts of a sample are grouped together in “Phylum <5% abund.”. The left panel labeled “N’ represents the birds as nestlings and the right panel labeled “SY” represents the birds as second-year individuals.

